# An Effective Method for Determining the Degree of Oligomerization of hnRNPA2 Low Complexity Domain

**DOI:** 10.1101/2025.02.24.639801

**Authors:** Paulina Żeliszewska, Zbigniew Adamczyk, Pooja Shah, Anna Kluza, Aneta Michna, Anna Bratek-Skicki

## Abstract

Theoretical calculations and various experimental techniques were applied to determine fundamental physicochemical characteristics of the RNA-binding protein low complexity domain (hnRNPA2 LCD), in sodium chloride solutions. The protein monomer size, cross-section area, the dependence of the nominal charge on pH, and its isoelectric point were predicted. These theoretical data allowed one to analyze and interpret the adsorption of hnRNPA2 LCD molecules on mica, which was investigated by the streaming potential technique, and on polymer particles, acquired by laser Doppler velocimetry. It was shown that the protein adsorbed in the form of oligomers whose size was resolved by atomic force microscopy. In the case of the adsorption on particles, the oligomer size and zeta potential were derived by applying the general electrokinetic model. Additionally, the electrokinetic properties of the hnRNPA2 LCD functionalized particles were determined and compared with the bulk protein properties. Using these results, a fast and easy method for quantifying the oligomerization kinetic of unstable protein solutions was developed.

## 1. Introduction

hnRNPA2/B1 is one of the RNA-binding proteins and is widely expressed in various tissues. It plays an important role in RNA stabilization, trafficking, translation, and splicing [1-4]. The hnRNPA/B protein group consists of A1, A2/B1, A3, and A0, forms which are structurally conserved and functionally complementing groups of proteins[5-8] involved in many important cellular functions. Previous reports presented that A2 and B1 have a little different affinity towards RNA binding regions [9-11] suggesting that they may be responsible for different functions. However, most reports on hnRNPA2/B1 either have not proved the difference between the two subtypes or were studying the main type A2 that is present in most tissues[12-15]. Therefore, this paper particularly focuses on hnRNPA2. One of its roles is the protection of mRNAs, an element of stress granules, one of the membrane-less organelles (MLOs) [1-4, 16]. It should be mentioned that hnRNPA2 is linked not only to numerous biological processes but also to diseases, particularly neurodegenerative disorders, e.g., mutations in the low complexity domain (LCD) in hnRNPA2 are responsible for multisystem proteinopathy including the development of amyotrophic lateral sclerosis (ALS), through promoting excess of hnRNPA2 into stress granules and driving the formation of cytoplasmic inclusions as reported in Ref.[2]. hnRNPA2/B1 is also involved in pluripotency, and hESC self-renewal [17]. The protein consists of 341 amino acids with two RNA-binding domains (RBD) and two RNA recognition domains (RRMs). In the end, the protein has a 161-residue C-terminal low-complexity domain (LCD) which contains numerous Gly, Phe, Asn, Tyr, and Asp residues but a low number of small hydrophobic amino acids such as Val, Leu, and Ala where Ile is absent. These regions are responsible for transient interactions between the numerous proteins involved not only in the stress granule formation but also in other MLOs [1, 18-20].

It has been shown that under cellular stress aggregated hnRNPA2 is present in cytoplasmic inclusions [2]. Similar results were obtained in vitro where under specific conditions the protein first formed liquid droplets, then hydrogel, similar to those observed in MLOs, and later fibrils [1, 18, 21, 22]. The formation of a dense network is possible by a process named liquid-liquid phase separation (LLPS) in a saturated solution of macromolecules [23-26].Phase transitions such as LLPS are initiated by the nucleation phenomenon, a thermodynamic process described, for the first time, by Gibbs in Ref.[27, 28].The nucleation process has been studied in a wide range of scientific fields[29]. During the past decades, there has been a renewed interest in developing the theory, which has resulted in new kinetic models of proteins forming aggregates linked to many neurogenerative diseases, e.g., Alzheimer’s Amyotrophic Lateral Sclerosis, and Parkinson’s disease [30-36]. One of the new developments in the field was work recently presented by Martin et al.[36]. The authors characterized phase boundaries and the size distributions of oligomers of hnRNPA1 LCD before its phase separation using rapid-mixing time-resolved small-angle X-ray scattering (SAXS) approaches. However, this issue has not been adequately studied and investigated for the hnRNPA2 LCD. This was caused by the lack of essential information about the hnRNPA2 subunit, particularly its effective charge in electrolyte solutions at various pHs and its isoelectric point, which controls the protein oligomerization processes.

Therefore, the main goal of this work was to determine the essential physicochemical characteristic of hnRNPA2 molecules in electrolyte solutions by applying theoretical calculations and thorough measurements of its adsorption on different solid/electrolyte interfaces comprising mica and polymer microparticles leading to the protein corona formation. The amino acid sequence of the studied domain is presented in Fig.1S in the Supplementary Material section. In this way, a comprehensive insight into the protein oligomerization mechanism is gained, which is not feasible regarding classical bulk measurements such as static or dynamic light scattering. In particular, the oligomerization number and their structure can be determined via the *in situ* laser Doppler velocimetry measurements interpreted using the electrokinetic model derived in [37].

Except for the significance to basic sciences the newly acquired results enable to a develop a robust method for quantifying oligomerization kinetic of unstable hnRNPA2 LCD protein solutions.

## 2. Materials and Methods

The plasmid hnRNPA2_LCD_WT encoding hnRNPA2 (Uniprot ID: P22626) subunit was gifted by Prof. N.L. Fawzi (Brown University, Providence, RI, USA). The plasmid is available at Addgene under catalog number #98657 and enables expression of 6xHis-tagged hnRNPA2 residues 190-341 (6xHis-hnRNPA2_LCD) with Tobacco Etch Virus (TEV) protease cleavage site between 6xHis-tag and hnRNPA2 subunit [20].

Freshly cleaved mica sheets (Continental Trade, Poland) were used for the hnRNPA2 LCD adsorption using atomic force microscopy and the streaming potential method. The thin sheets of mica were freshly cleaved before each experiment.

Polystyrene particles with sulfonate, negatively charged groups (PS800),used for hnRNPA2 LCD adsorption, were synthesized by the Goodwin protocol [38].

Sodium chloride (NaCl) and phosphate buffered saline (PBS) were purchased from Avantor Performance Materials Poland S.A. and Biomed (Lublin, Poland), while hydrochloric acid (HCl) was the product of Merck KGaA company Germany). The used salts, acid and buffers were used without further purification. Deionized water was prepared using the Milli-Q Elix & Simplicity 185 purification equipment purchased from Millipore SAS Molsheim, France.

The temperature of all experiments was constant equal to 298 K.

### Protein production

Plasmid hnRNPA2 LC WT (Addgene, cat. no. #98657) was transformed into Escherichia coli One Shot BL21 Star cells by heat-shock method [39]. A single colony was picked up and their growth lasted one night in an LB medium containing 50 μg/mL kanamycin. One liter of terrific broth media containing 50 μg/mL kanamycin was inoculated with the preculture, and the cells were grown at 37°C, 180 rpm until an OD600 reached 0.6 – 0.8. The temperature was decreased to 26°C and the protein expression was initiated with 1 mM isopropyl β-D-1-thiogalactopyranoside (IPTG). After an overnight incubation (26°C, 150 rpm) the cells were harvested by centrifugation (4000 rpm, 60 minutes, 4°C). The cells were stored at -80°C until further use. Following thawing, cells were resuspended in lysis buffer: 20 mM Tris-HCl, 500 mM NaCl, 10 mM imidazole, pH 8.0 at 4°C supplemented with 1 mM DTT, 0.1 mM phenylmethylsulfonyl fluoride, 0.5 mM benzamidine hydrochloride, and 1 tablet of Roche complete EDTA-free protease inhibitor per 50 mL. The cells were lysed by sonication on ice (5 s pulse on, 5 s pulse off, 60% amplitude; Sonopuls, Bandelin). Inclusion bodies containing 6xHis-hnRNPA2_LCD were centrifuged at 40 000 g for an hour at 4°C. Pellet was resuspended in 20 mM TRIS-HCl buffer, 500 mM NaCl, 10 mM imidazole, 1 mM DTT, 3 mM urea, pH 8.0, and the suspension was once again sonicated on ice (10 s pulse on, 10 s pulse off, 70% amplitude). The cell debris was then removed by centrifugation for one hour at 20,000 g at 4 °C. The supernatant was filtered through a 0.45-μm pore syringe filter and applied to the IMAC column (Nuvia IMAC Column, Ni-charged, 5 mL) connected to Bio-Rad NGC Chromatography System. The contaminants were washed (20 mM TRIS-HCl, 500 mM NaCl, 25 mM imidazole, 1 mM DTT, 3 M Urea, pH 8 at 4°C), and 6xHis-hnRNPA2 LCD eluted with gradient elution up to 500 mM imidazole. Buffer was exchanged to TEV cleavage buffer (50 mM sodium dihydrogen phosphate, 200 mM sodium chloride, 3M urea, pH 7.0) on desalting column HiPrep 26/10 Desalting (Cytiva). 6xHis-tagged TEV protease was added to 6xHis-hnRNPA2 LCD in 1:10 molar ratio to remove the 6xHis-tag. Following an overnight incubation period at room temperature, the solution was applied to the IMAC column to remove the cleaved tag and TEV protease. The flowthrough containing the hnRNPA2 LCD was collected and loaded on HiLoad 16/600 Superdex 200 pg (GE Healthcare) previously equilibrated with 50 mM sodium dihydrogen phosphate, 200 mM sodium chloride, 3 M urea, pH 7. The hnRNPA2 LCD was dialyzed at 4°C into 10 mM citric acid, pH 3.5. The protein was aliquoted and stored at -80°C until further use. The concentration of hnRNPA2 LCD was estimated by absorbance at 280 nm based on the extinction coefficient calculated by protparam (cit https://web.expasy.org/protparam/protpar-ref.html). The purity of the sample was estimated based on SDS-PAGE stained with BlueStain Sensitive Plus (EURx)and presented in Fig.2S.

### 2.2 Experimental Methods

#### 2.2.1 Atomic Force Microscopy Measurements

Freshly cleaved mica sheets were used in protein adsorption experiments carried out under diffusion-controlled transport in a thermostated cell.

Three mica sheets/per one protein concentration were vertically immersed into the hnRNPA2 LCD solution, c = 0.5 mg/L, 10 mM NaCl, pH 3.5, and incubated for 15 min. Afterward, the hnRNPA2 LCD-covered mica sheets were rinsed a few times (usually for 30 seconds) using an electrolyte solution of the same pH and ionic strength that was used for the hnRNPA2 LCD adsorption experiment. The mica sheets with the adsorbed hnRNPA2 LCD molecules were observed by atomic force microscopy. Micrographs of the adsorbed protein layers were acquired by ambient air AFM imaging using the NT-MDT Solver device with the SMENA - B scanning head. The measurements were performed using semi-contact mode and polysilicon cantilevers NSG-03 with resonance frequencies of 47-150 kHz, and a tip radius of 10 nm. The images of adsorbed protein molecules were acquired within the scan range of 2.0 μm per 2.0 μm, over 10-20 randomly chosen areas of the mica sheet, which ensures a relative precision of the coverage determination of about 3%. All the images were flattened using an algorithm provided by the instrument producer.

#### 2.2.3 Bulk Characteristics of hnRNPA2 LCD protein and PShnRNPA2LCD protein corona

Bulk characteristics of the hnRNPA2 LCD protein, latex particles, and latex particles with adsorbed hnRNPA2 LCD were acquired using the dynamic light scattering (DLS) that yielded the diffusion coefficient, and the laser Doppler velocimetry (LDV) furnishing to measure the electrophoretic mobility, (Zetasizer Nano, Malvern Instruments, United Kingdom).

The electrophoretic mobility measurements of hnRNPA2 LCD were performed in the range of pH between 4.0 and 11.0 at *I* =10^−3^M, c = 1000 mg/L. The hnRNPA2 LCD adsorption on latex particles was measured by recording the changes in its electrophoretic mobility (zeta potential) induced by this process. The experimental protocol consisted of the following steps: (i) measuring the electrophoretic mobility of bare latex particles in suspensions of the concentration of 100 mg/L; (ii) formation of hnRNPA2 LCD monolayers on latex particles by mixing the hnRNPA2 LCD solution of an appropriate concentration ranging from 0.1 to 4.0 mg/L with the latex suspension having a concentration of 100 mg/L. The stock concentration of hnRNPA2 LCD was 125 mg/L. The adsorption time of the protein on PS800 was 600 s.

Since the electrophoretic mobility measurements were a significant parameter for obtaining reliable data, a special care was applied to ensure the accuracy of he obtained data. Thus, for every experimental point (defined by various pHs and/or hnRNPA2 LCD concentration) up to ten independent measurements of the electrophoretic mobility were carried out. Moreover, each of these measurements was repeated three times using a fresh sample.

The experimental data was used for calculation of the hydrodynamic diameter of the microparticles using the Stokes-Einstein equation and the zeta potential using the Smoluchowski-Henry relationship.

#### 2.2.3 Streaming Potential Measurements

The streaming potential experiments were conducted in a parallel-plate channel created using two mica sheets, separated by a Teflon gasket, following the procedure described in previous papers [40]. Measurements were taken *in situ* using a pair of reversible Ag/AgCl electrodes. Several measurements were performed at four different pressure differences that drove the flow through the cell. This allowed us to obtain the slope of the streaming potential vs. hydrostatic pressure difference dependence, which was used to calculate the zeta potential of both the bare and protein-covered mica substrate using the Smoluchowski equation [41].

Protein adsorption was conducted within the streaming potential cell, specifically in the channel designated for SPM. During the adsorption process, the protein solution flowed through the cell channel at a constant volumetric rate of 0.02 mL min^-1^ for the desired adsorption time (0-100 min) at *I* = 10 mM, and the pH range from 4.0 to 9.0.

## 3. Results and Discussion

### 3.1 Theoretical Calculations of Molecule Size and Charge

The primary structure (amino acid sequence) of the hnRNPA2 LCD monomer was previously given in Ref.[23]. Using these data, the molar mass, denoted as *M*1, was calculated to be 14,000 g mol^-1^ (14 kDa). It is assumed that the molecule density is equal to that of the lysozyme molecule, that is 1.35 g cm^-3^ given its similar molar mass of 14,300 g mol^-1^ [42]. Using this value one can predict that the volume of the hnRNPA2 LCD monomer is equal to 17.2 nm^3^. Consequently, the diameter of the equivalent sphere

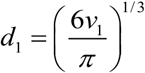

is equal to 3.2 nm. Additionally, the radius of gyration, the cross-section area, and the diffusion coefficient of the monomer can be calculated and compared with those pertinent to the lysozyme molecule, see Table 1.

**Table 1.**
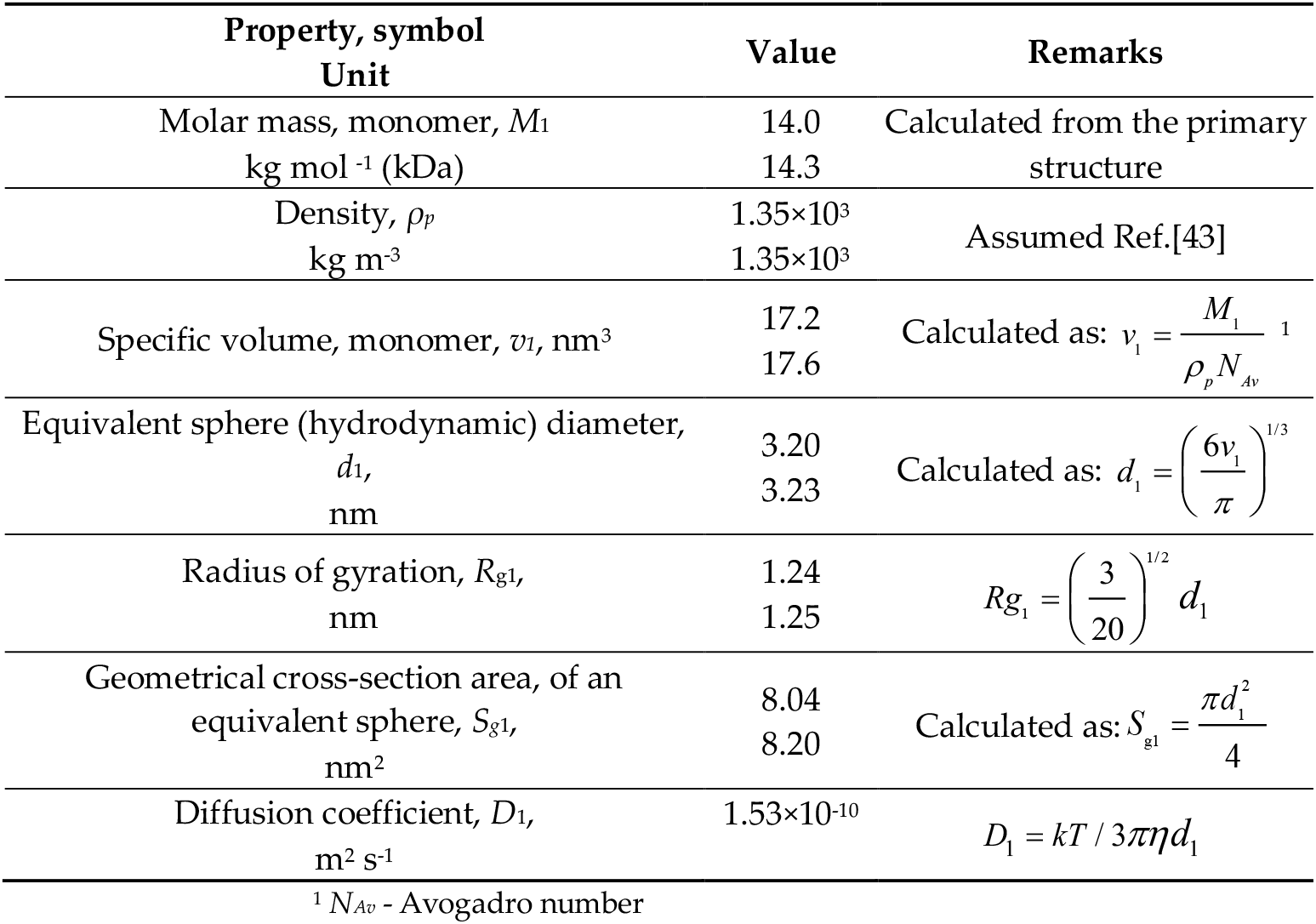
Predicted parameters of hnRNPA2 LCD subunit monomer (upper numbers) and the lysozyme molecule (lower numbers).

The parameters in Table 1 were obtained assuming a quasi-spherical shape of the hnRNPA2 LCD monomer, providing a useful first-order approximation for interpreting protein adsorption experiments. This approach is especially relevant since the crystallographic structure of the molecule has not yet been determined, making it impossible to derive a more precise shape from the molecular dynamic modeling. Notably, it was shown in Refs.[44, 45] that the disordered (unfolded) protein molecule shape in good solvents can well be approximated as an excluded volume random coil exhibiting a quasi-spherical shape. Quasi-spherical structures of hnRNPA2 LCD have been also obtained using the AlphaFold algorithm[46-50] and are presented in Fig.3Sa.

The solvent-accessible (nominal) charge of the hnRNPA2 LCD molecule was calculated using the PROPKA algorithm [51-53]. The results are shown in Figure. 1, indicate that the hnRNPA2 LCD carries a charge of 8*e* at pH 4, remaining positive up to pH 9. A similar behavior exhibits the charge vs. pH dependence predicted for the lysozyme molecule, although its charge was markedly larger for the entire range of pH and equal to 15*e* at pH four.

**Figure 1.**
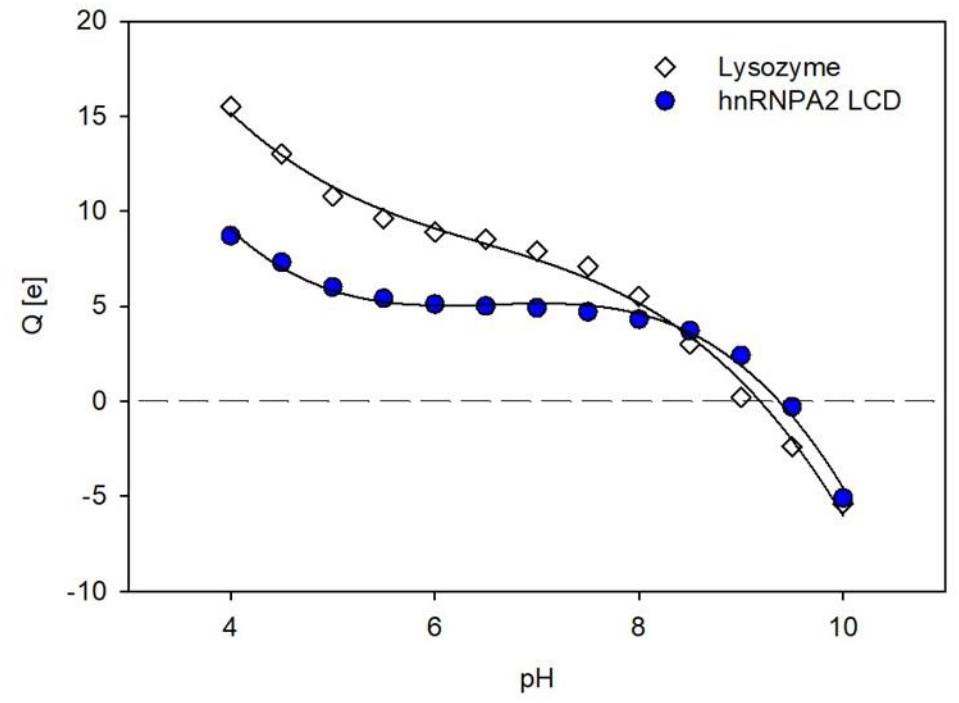
The nominal charge (*Q*) of the chicken-lysozyme molecule (◊) and the hnRNPA2 LCD monomer 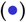versus pH calculated using the PROPKA 3.0 algorithm.

### 3.2 Experimental Results

#### 3.2.1 Characteristics of Microparticles and the Substrate used in Adsorption Experiments

The hydrodynamic diameter of the polystyrene microparticles (hereafter referred to as PS800) determined via the DLS measurements was equal to 820 ± 20 and 810 ± 20 nm at NaCl concentrations of 10 and 150 mM, respectively, for the pH range of 4 to 9. The zeta potential of the particles determined via the LDV electrophoretic mobility measurements was -97 and -105 mV at pH 4 and 7.4, respectively (for 10 mM NaCl). The pH dependence of the zeta potential of the PS800 particles is shown in Figure 2, part (a).

**Figure 2.**
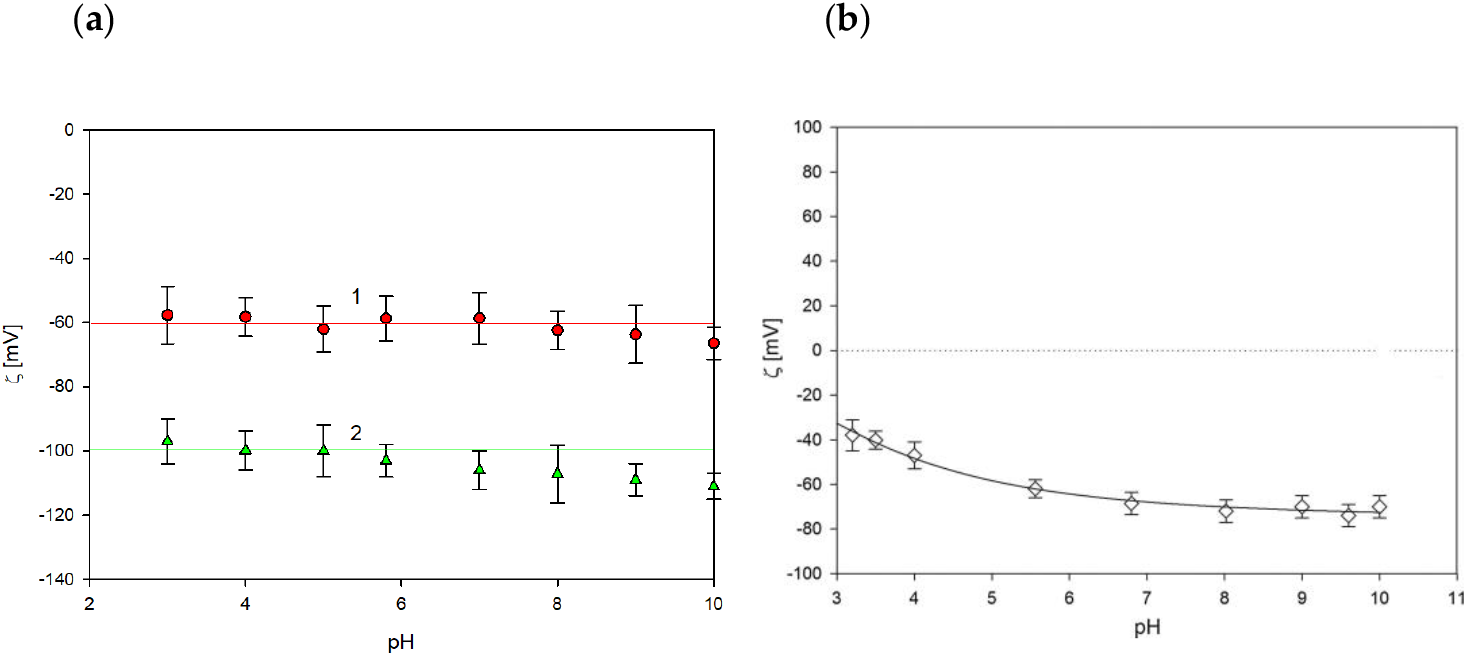
(a) Dependence of the zeta potential of the polymer microparticles PS800 on pH determined by the LDV method: 1. 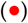 150 mM NaCl, 2. 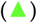 10 mM NaCl; (b) Dependence of the zeta potential of mica on pH determined by the SPM in 10 mM (◊). Reprinted from[54] Copyright © 2014 Elsevier Inc., with permission from Elsevier. The solid lines are guides for the eyes.

Physicochemical characteristics of the mica substrate, including surface topography and the zeta potential, were acquired by AFM and SPM as described above. The root mean square (rms) roughness of the mica sheets used in the AFM and SPM was below 0.1 nm. The zeta potential of mica was negative for the entire pH range and at 10 mM NaCl concentration, consistent with literature data [54] (see Figure 2 part b).

#### 3.2.2 hnRNPA2 LCD Molecule Characteristics: AFM and SPM

Primarily, the electrophoretic mobility of the hnRNA2 LCD molecules in NaCl solutions was measured by the LDV method and the zeta potential was calculated using the Henry formula. The results presented in Figure 3 indicate the mobility was positive for pH up to 8 and then abruptly decreased attaining a large negative value. Interestingly, this trend exhibits a qualitative agreement with the nominal charge vs. pH dependence shown in Figure 1. However, the zeta potential of the molecules was rather low equal to 11 mV at pH 4 and 1.6 mV at pH 7.4. The experimental data shown in Figure 3 were used in the last section of this work to calculate the electrokinetic (compensated) charge of the hnRNPA2 LCD molecules.

**Figure 3.**
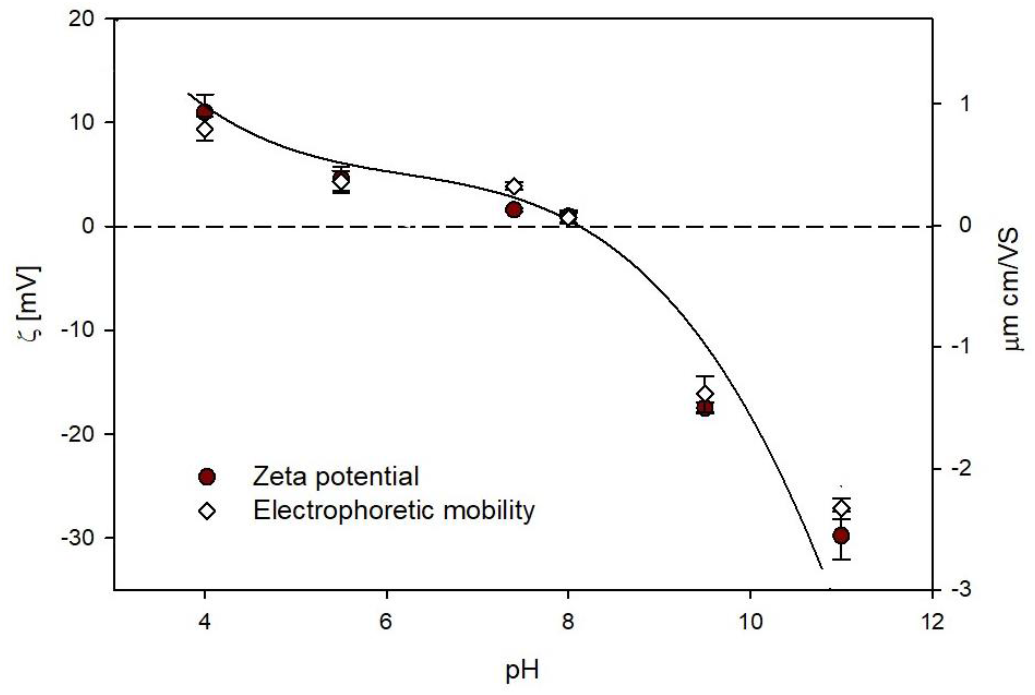
Dependence of the zeta potential 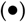 (left hand axis) and the electrophoretic mobility (◊) of the hnRNA2 LCD molecules on pH determined by the LDV method in 10 mM NaCl. The line is the guide for the eyes.

On the other hand, the size distribution of hnRNPA2 LCD oligomers in NaCl solutions was determined according to the previously applied procedure[55, 56] via controlled adsorption at mica under diffusion conditions. These measurements, conducted with a protein solution of ca 1 mg L^-1^, offer advantages compared to the DLS method, which requires concentrations of around 300 mg L^-1^ to produce reliable results. In Figure 4 and Fig.6S, the AFM micrographs of the protein layers acquired for the adsorption time of 15 min at pH 3.5 and 10 mM NaCl are shown for (a) freshly prepared protein solution (the bulk concentration of the stock suspension 125 mg L^-1^) and (b) solution prepared from protein stored over 12 months at the same bulk concentration.

**Figure 4.**
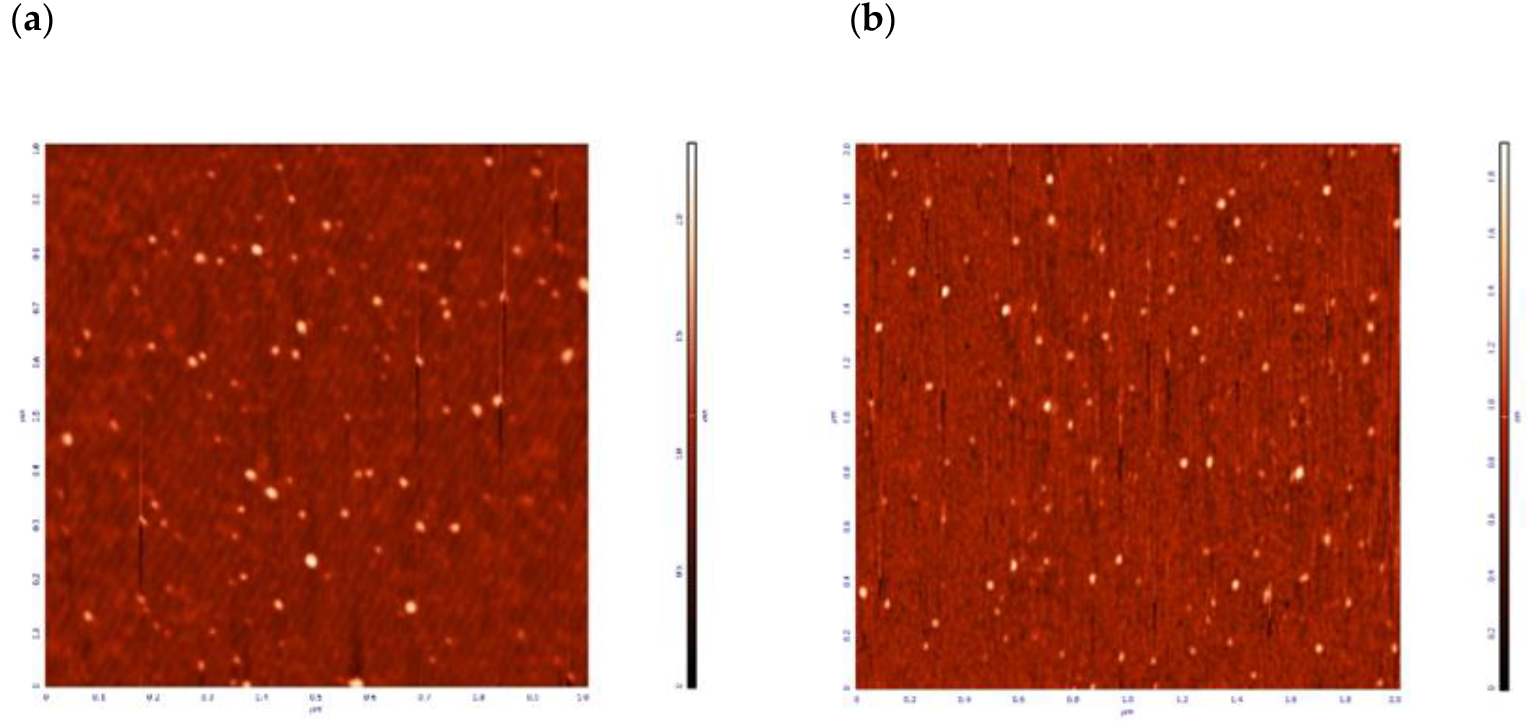
AFM micrographs of hnRNPA LCD protein layers on mica after 15 minutes of adsorption. Experimental conditions: bulk protein concentration *c*_*b*_ = 0.5 mg L^-1^, pH 3.5, 10 mM NaCl. (**a**) Freshly prepared protein solution; (**b**) Solution prepared from protein stored for over 12 months.

The size distribution of oligomers was determined by measuring their dimensions in two perpendicular directions and taking the average value. The resulting size histograms, created from measurements of approximately 100 individual oligomers, are shown in Figure 5. For freshly prepared protein solution, the average oligomer size was 11 nm, while for the solution prepared from stored protein, the average oligomer size was markedly larger and equal to 21 nm. Thus, these AFM investigations showed that the oligomer sizes significantly exceeded the predicted hnRNPA2 LCD monomer size of 3.2 nm and that the oligomerization degree increased with storage time [50]. Height profile measurements of the oligomer cross-sections confirmed that, in both cases, their shape could be approximated as spherical. Additionally, simulations of oligomeric structures using the AlphaFold algorithm [46] also show quasi-spherical shapes of structures like dimers or 13-mers (see Fig.3S b and c). On the other hand, the information about the electrokinetic properties of the hnRNPA2 LCD layers adsorbed on mica was derived from the SPM carried out using the microfluidic channel cell, as described above.

**Figure 5.**
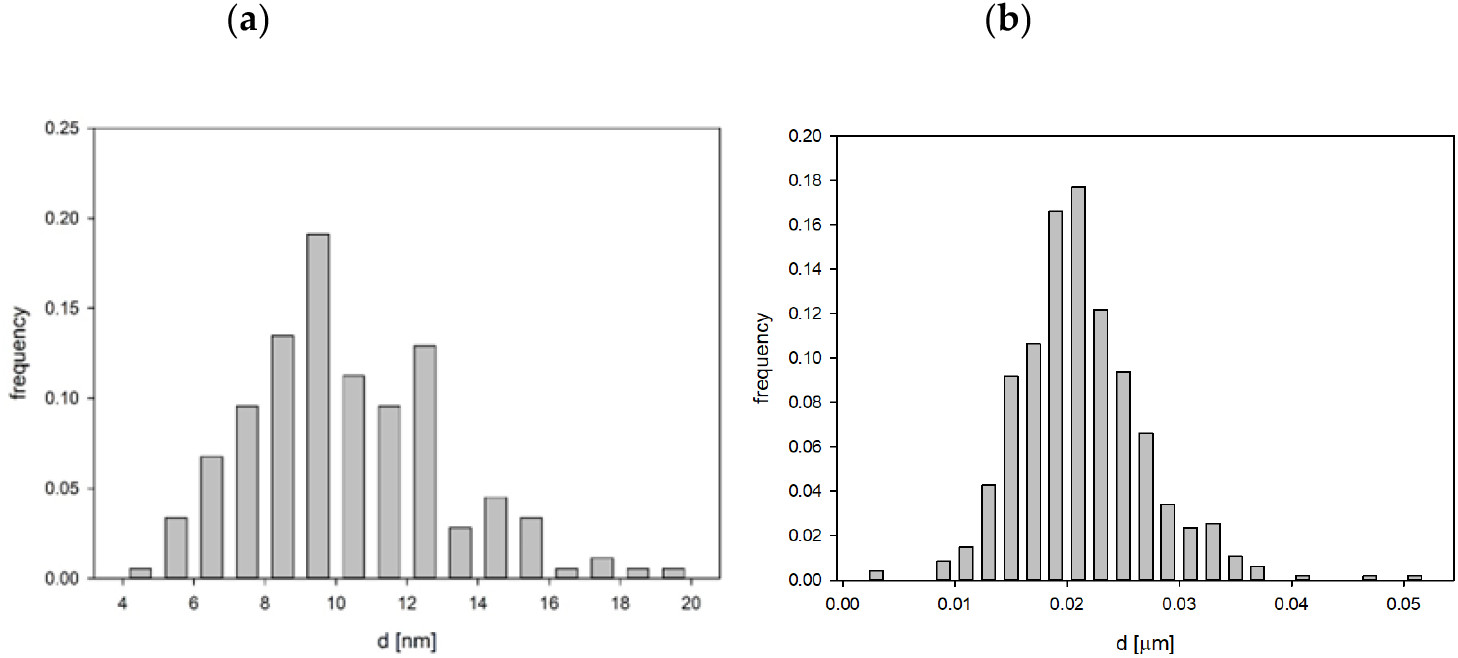
Histogram of hnRNPA2 LCD protein oligomers size distribution on mica (adsorption conditions as in Figure 4). (**a**) Freshly prepared protein solution, average oligomer size 11 ± 4 nm; (**b**) Solution prepared from protein stored for over 12 months at -70 C, average oligomer size 21 ± 5 nm.

Results showing the dependence of the zeta potential of mica on the adsorption time of hnRNPA2 LCD acquired at pH 4.0 and the protein bulk concentration of 2.5 mg L^-1^ (in 10 mM NaCl) are presented in Figure 4S. One can observe that the initially negative zeta potential of bare mica (equal to – 36 mV) monotonically increased with the adsorption time attaining a positive plateau value of 10 mV after 100 min. It is also worth mentioning that the consecutive desorption run where pure electrolyte was flushed through the cell induced a negligible change in the plateau zeta potential.

The experimental results were interpreted in terms of the convective-diffusion theory [55, 57] which yielded the protein’s dimensionless coverage as a function of the adsorption time. This dependence was subsequently transformed to express the dependence of the zeta potential on the adsorption time, using the electrokinetic model discussed in the next section. As shown in Figure 4S, the experimental data well agree with this theoretical model.

The pronounced stability of the hnRNPA2 LCD layer on mica enabled to determine its zeta potential via the streaming potential measurements. The pH was gradually increased from 4.0 to 9.0 by adding NaOH and then decreased back to pH 4.0 by adding HCl. The results of these measurements shown in Figure 5S, indicate that the initially positive zeta potential of the layer changes its sign at pH 4.5 and attains a negative value of -35 at pH 9.0. Interestingly, after the pH reversal, the zeta potential returns to the initial value of approximately 12 mV at pH 4.0, confirming its stability.

#### 3.2.3 hnRNPA2 LCD Adsorption at Polymer Microparticles

Compared to the above tedious experiments for planar substrates, investigations of protein adsorption at carrier microparticles exhibit pronounced advantages. Primarily, the monolayer formation time in the latter case is considerably shorter, typically of the order of a second and it does not depend on the bulk protein concentration. Therefore, experiments can be efficiently carried out even for the nanomolar range of protein solution concentration, which is especially advantageous for recombinant proteins acquired in small quantities. Additionally, the coverage of the protein layer formed on the carrier microparticles (often referred to as the protein corona) [58, 59]can be effectively monitored by the electrokinetic measurements carried out using the LDV method as described above.

The results showing the dependence of the zeta potential of PS800 microparticles on the initial concentration of hnRNPA2 LCD at pH 4.0, for 10.0 and 150 mM NaCl, are plotted in Figures 6 and 7, respectively. To assess the significance of oligomerization, data from both freshly prepared and stored protein solutions are included.

**Figure 6.**
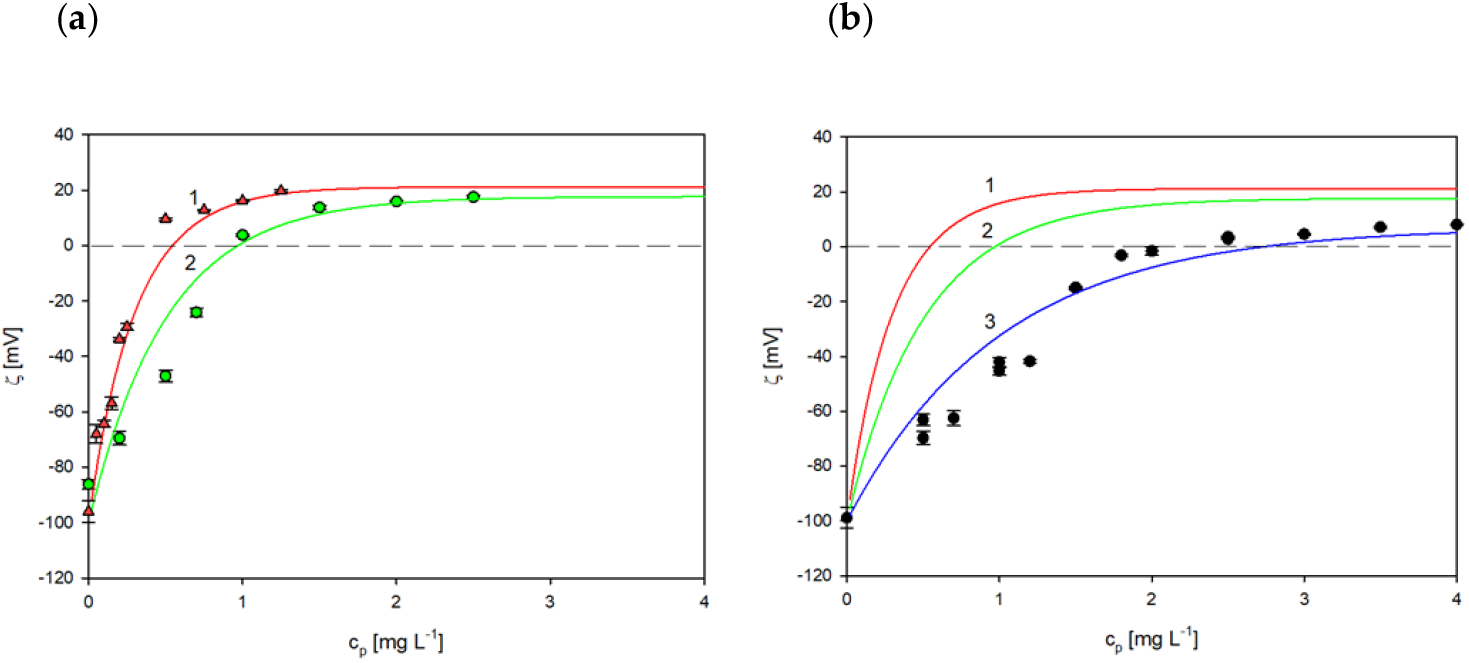
The dependence of the zeta potential of polymer microparticles (PS800) on the hnRNPa2 LCD concentration; adsorption conditions: protein volume 5 ml, particle suspension volume 5 ml, particle concentration 100 mg L^-1^, pH 4.0, 10 mM NaCl. (**a**) Freshly prepared protein solution (stock suspension concentration of 50 mg L^-1^); the triangle points show the experimental results obtained from the LDV measurements (stock suspension concentration of 50 mg L^-1^), the circles show the experimental points (stock suspension concentration of 125 mg L^-1^), the solid lines show the theoretical results calculated for adsorption of the monomer (red line 1) and oligomer composed of 13 molecules (green line 2);(**b**) Stored protein, the points denote experimental results obtained from the LDV measurements, the solid lines show theoretical results calculated for adsorption of the monomer (red line 1), oligomer composed of 13 molecules (green line 2), and the oligomer composed of 64 molecules (blue line 3).

**Figure 7.**
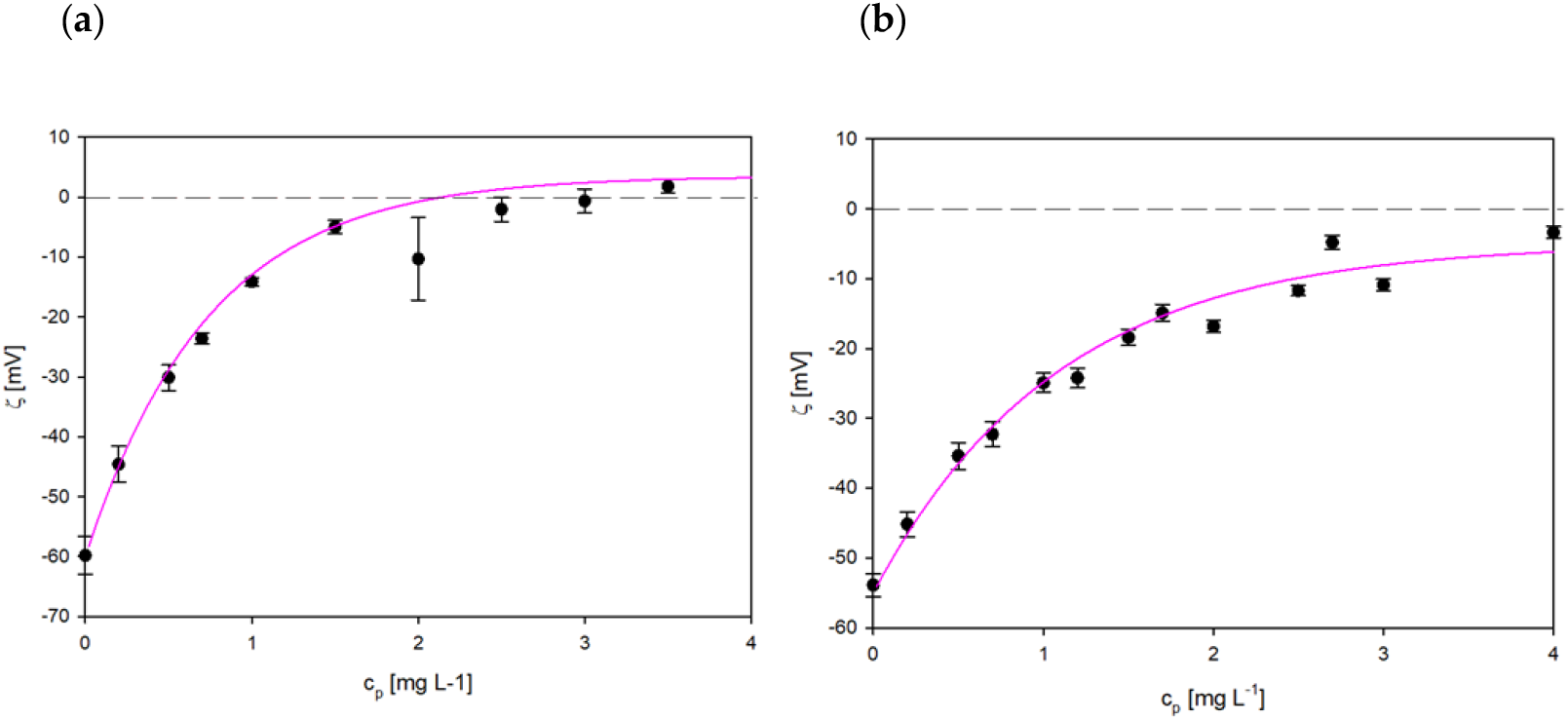
The dependence of the zeta potential of polymer microparticles (PS800) on the bulk hnRNPA2 LCD concentration, pH 4.0, 150 mM NaCl, (**a**) Freshly prepared protein solutions (stock concentration 125 mg L^-1^) the points denote experimental results obtained from the LDV measurements, the solid line shows the theoretical results calculated for adsorption of the oligomer composed of 64 molecules; (**b**) Solutions prepared from stored stock suspensions, the points denote experimental results obtained from the LDV measurements, the solid lines show the theoretical results calculated for adsorption of the oligomer composed of 110 molecules.

One can observe in Figure 6 that for 10 mM NaCl the initially negative zeta potential of polymer microparticles rapidly increased with the concentration of the protein and changed its sign to positive at *c*_*p*_ of 0.5 and 1.0 mg L^-1^ for solutions prepared from the stock suspensions with the concentration of solution concentration 50 and 125 mg L^-1^, respectively. This corresponds to the hnRNPA2 LCD monomer concentration of 35-70 nM. Given that the zeta potential can be determined with a relative precision of 5%, this result indicates that using the LDV method, the protein concentration can be determined for the nanomolar range. However, for the protein concentration larger than 1 mg L^-1^ the increase in the zeta potential was less abrupt, and finally, a plateau value of 20 mV was attained in both cases. The increase in the zeta potential for the stored protein was less rapid and its plateau value was about 5 mV.

Analogous dependencies acquired for the 150 mM NaCl solution are shown in Figure 7. Both for the freshly prepared and stored protein solutions the increase in the particle zeta potential was less abrupt compared to 10 mM NaCl and the plateau values were close to zero.

To quantitatively interpret these results, the mass coverage of the protein layer on polymer microparticles *Γ* should be known. Assuming irreversible adsorption, the calculation can be performed based on mass balance using the following formula

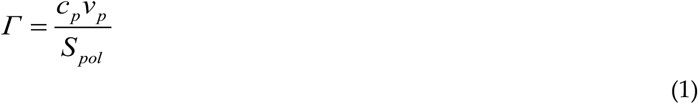

where *c*_*p*_ is the protein concentration in the solution (before mixing), *ν*_*p*_ is the solution volume and *S*_*pol*_ is the surface area of the microparticles given by

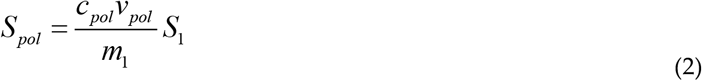

where *c*_*pol*_ is the particle suspension concentration, *ν*_*pol*_ is the particle volume (before mixing), and *S*_1_, *m*_1_ is the surface area and mass of a single polymer particle, respectively. For spherical particles one has

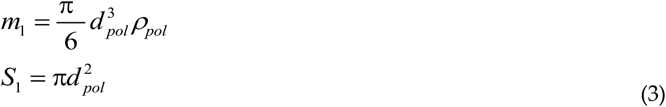

where *d*_*pol*_, *ρ*_*pol*_ are the polymer particle diameter and the density, respectively.

Considering equations 2-3 one obtains the following formula for the mass average of the protein on the polymer microparticles

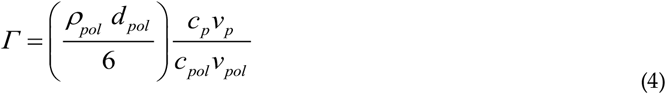

Consequently, the dimensionless coverage of protein molecules is given by

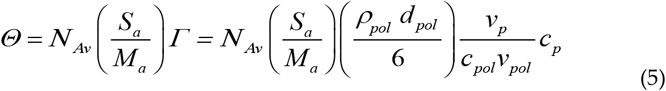

where *N*_*Av*_ is the Avogadro constant, and *S*_*a*_ is the characteristic cross-section area of the molecule (oligomer) and *M*_*a*_ is the molecule (oligomer) molar mass.

The oligomer molar mass can be calculated as *n*_*a*_ *M*_*1*_ where *n*_*a*_ is the oligomerization number and its cross-section area was calculated for *n*_*a*_ *>* 10 from the dependence *S*_*a*_ = *C*_*H*_ *n*_*a*_^*2/3*^ *S*_*1*_ where the *C*_*H*_ constant was 1.22 assuming a random structure of the oligomers. Upon defining the dimensionless coverage one can interpret the experimental results in terms of the general electrokinetic model [60] where the following expression for the zeta potential of the protein layer (corona) *ζ*(*Θ*) on carrier microparticles was formulated

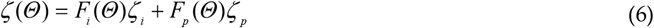

where *ζ*_*i*_ is the bulk zeta potential of the polymer microparticles, *ζ*_*p*_ is the bulk protein (oligomer) zeta potential, and *F*_*i*_ (*Θ*), *F*_*p*_ (*Θ*) are the dimensionless functions. The *F*_*i*_ function describes the damping of the local flow by the adsorbed molecule layer and the *F*_*p*_ function characterizes the contribution to the zeta potential stemming from the electric double-layers around adsorbed molecules.

The results calculated from equation 6 for the hnRNPA2 LCD monomer and oligomers composed of a various number of single molecules are plotted as solid lines in Figures 5S and 6. One can observe in the former Figure that for the experimental results acquired for the freshly prepared protein solutions (from the stock suspension of 50 mg L ^-1^) the experimental results are adequately described by the theoretical model pertinent to the hnRNPA2 LCD monomer. However, for the protein solutions prepared from the stock suspension of 125 mg L^-1^ the experimental results are adequately described by the theoretical model assuming the oligomerization degree of 13, which corresponds to its molar mass of 182 kg mol^-1^. In the case of stored stock solutions, the predicted oligomerization number was much larger, equal to 64, with a molar mass of 896 kDa.

The results acquired for 150 mM NaCl (Figure 7) indicate that the oligomerization degree of the freshly prepared protein solutions (stock suspension of 125 mg L^-1^) increased with the ionic strength. One can also argue that these results confirmed that by applying the controlled adsorption procedure one can prepare functionalized polymer microparticles (hereafter referred to as PShnRNPA2 LCD) with a controlled coverage of hnRNPA2 LCD and the zeta potential. Additionally, this method, requiring nM quantities of protein, can be used for sensitive estimation of the oligomerization state of protein solutions. The stability of the PShnRNPA2 LCD particles upon storage was investigated in a series of experiments where their hydrodynamic diameter and zeta potential were determined as a function of time. The results shown in Figure 8 indicate that their hydrodynamic diameter did not change for the storage time up to 60 hours, whereas the zeta potential remained stable for the time up to 30 hours. These facts confirm that the electrokinetic properties of the microparticles can be acquired in a facile way by applying the LDV measurements. Considering the large stability of the functionalized PShnRNPA2 LCD particles, one can perform their thorough electrokinetic characteristics. This is particularly significant considering that the corona formation only requires protein nM protein quantities, whereas the bulk LDV measurement (see Figure 3) requires at least two orders of magnitude larger protein quantities.

**Figure 8.**
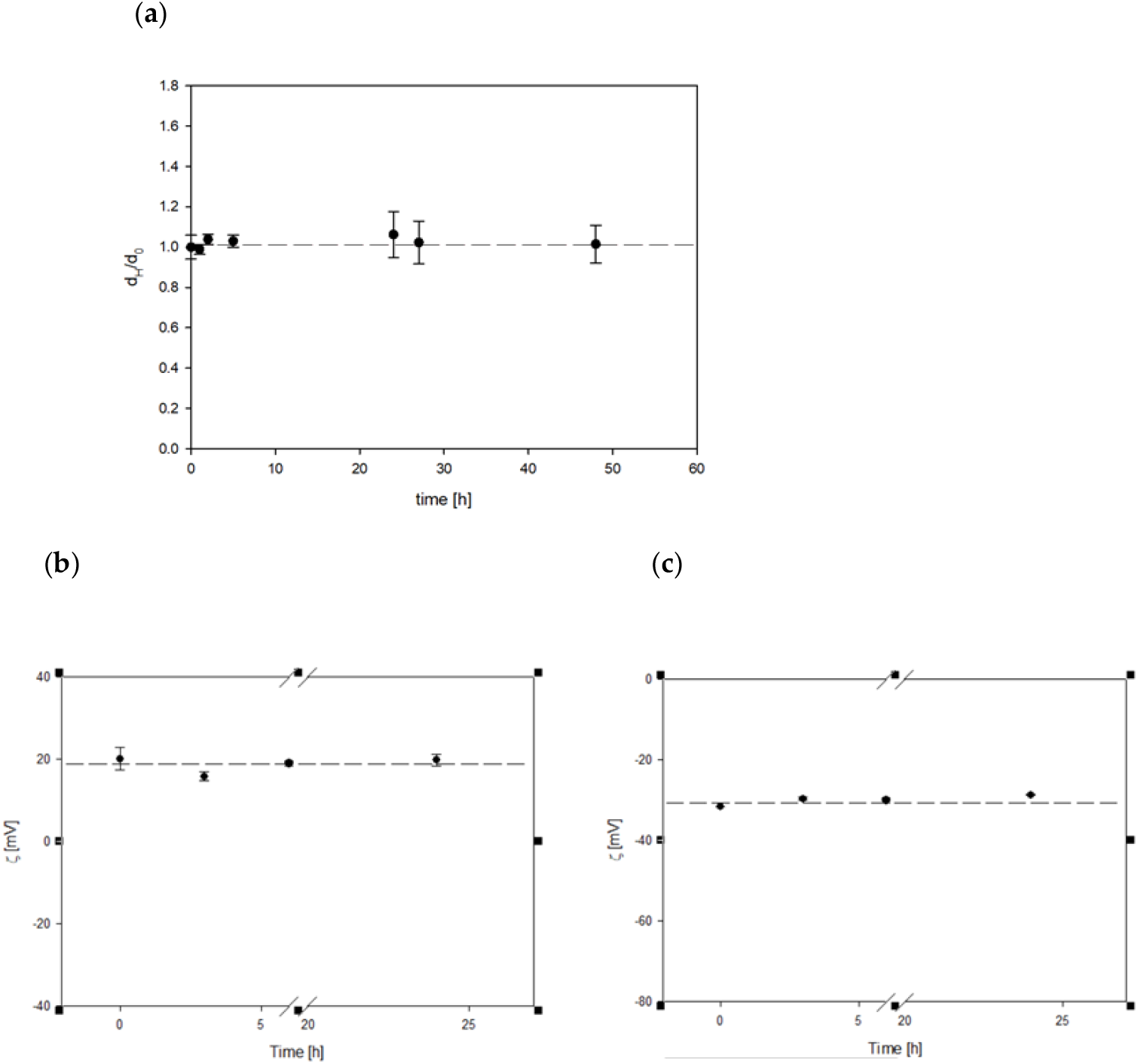
Stability of hnRNPA2 LCD functionalized particles determined by the DLS measurements. (**a**) pH 4.0, 10 mM NaCl, and by the electrophoretic mobility measurements; (**b**) pH 4.0, 10 mM NaCl; (**c**) pH 7.4, 10 mM NaCl. Such an electrokinetic characteristic expressed in terms of the dependence of the PShnRNPA2 LCD particle zeta potential on pH is shown in Figure 9. It can be seen that the zeta potential remained positive for pH up to 6 and for larger pHs attained large negative values. Therefore, the results obtained for the functionalized particles resemble those shown in Figure 3 acquired by applying the tedious LDV measurements for protein solutions.

**Figure 9.**
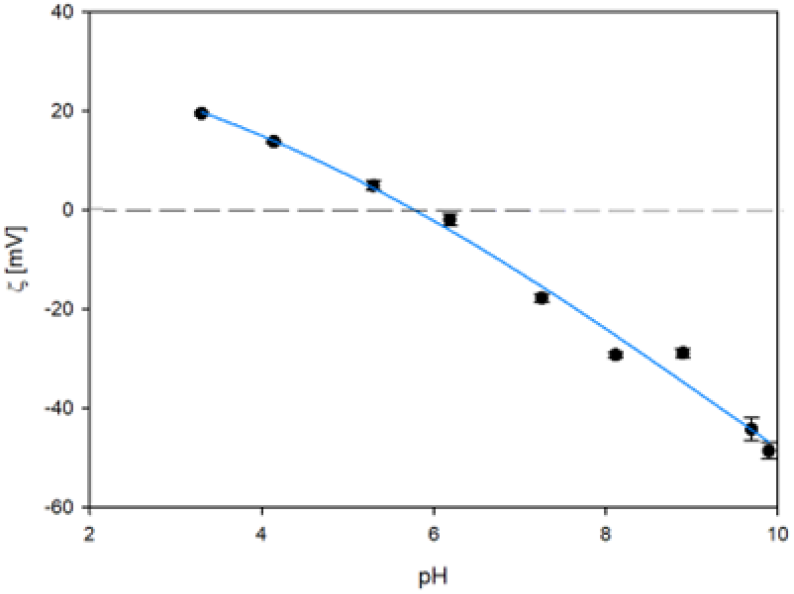
Dependence of the zeta potential of the PShnRNPA2 LCD particles on pH. The points represent experimental data acquired using the LDV method for 10 mM NaCl of freshly prepared solutions (stock suspension concentration 125 mg L ^-1^). The solid line is the guide for the eyes.

Additionally, using the dependence of the functionalized particle zeta potential on pH one can calculate the zeta potential of the protein oligomers in the bulk transforming equation 6 to the following form

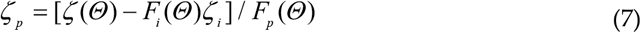

The results obtained considering that for protein coverage above 0.3, the *F*_*i*_*(Θ*) function exponentially vanishes and the *F*_*p*_(*Θ*) function approaches 0.7 are collected in Table 2. Additionally, using the zeta potential data one can calculate the electrokinetic charge of the hnRNPA2 LCD molecules using the following dependence

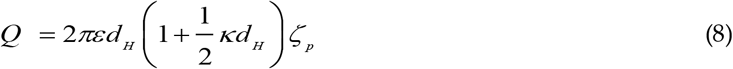

where *ε* is the electric permittivity of the electrolyte, *d*_*H*_ is the hydrodynamic diameter of the protein molecule (oligomer) and *κ*^−1^ is the electric double-layer thickness equal to 3.1 nm for the 10 mM NaCl solution.

**Table 2.** The zeta potential and the electrokinetic charge of the hnRNA2 LCD molecules vs pH derived from the LDV measurements for hnRNPA2 LCD protein solutions in the bulk and for the functionalized particles with the PShnRNPA2 LCD corona. The last column presents the nominal charge theoretically predicted (shown in Figure 1).

| pH | $\zeta_p$ [mV]<br>bulk | Q [e]<br>Bulk | $\zeta_p$ [mV]<br>corona | Q [e]<br>corona | Q [e]<br>theory |
| --- | --- | --- | --- | --- | --- |
| 3.5 | - | - | 33 | 4.2 | 10 |
| 4.0 | 11 | 1.4 | 24 | 3.1 | 8.0 |
| 5.5 | 4.6 | 0.6 | - | - | 5.0 |
| 6.0 | - | - | 4.4 | 0.57 | 5.0 |
| 7.4 | 4.1 (1.6) | 0.2 | -24 | -3.1 | 4.0 |
| 8.0 | 0.95 | 0.12 | -38 | -4.9 | 3.8 |
| 9.5 | -17 | -2.2 | -67 | -8.7 | 0 |
| 10 | -30 | -3.9 | - | - | - 5.0 |

Analyzing the results presented in Table 2 one can deduce that the electrokinetic charges derived from the bulk electrophoretic mobility measurements are significantly smaller than the nominal charge theoretically predicted using the Propka software (last column). This effect can be attributed to a significant compensation of the charge by counterions, the effect that was previously observed for other proteins such as the SARS-CoV -2 spike protein subunit and myoglobin [61]. It is also interesting to observe that the electrokinetic charge of the molecules derived from the LDV measurements for the functionalized particles was negative for pH below 6.0, where in contrast, the theoretically predicted value for the monomer was positive. This difference can be probably attributed to the polarization of the adsorbed hnRNPA2 LCD molecules by the strong electric field of the polymer particles bearing a negative charge for the entire pH range.

## 4. Conclusions

AFM investigations of the hnRNPA2 LCD molecule adsorption on mica under diffusion-controlled transport furnished reliable information about the stability of its solutions, particularly about the oligomerization degree. It was confirmed that the protein layer consisted of quasi-spherical oligomers with the size depending on the storage time. It was also shown that the oligomerization degree increased with the ionic strength of the electrolyte suggesting a significant role of the electrostatic interactions.

The oligomerization mechanism of the protein was also effectively investigated by applying the adsorption at carrier particles where the monolayer formation time was considerably shorter than for planar substrates. Additionally, the precision of these measurements was larger, therefore reliable results could be acquired for nanomolar amounts of the protein. These experimental results were quantitatively interpreted in terms of the electrokinetic model that yielded valid information about the protein oligomerization degree and its surface coverage. This knowledge can be exploited to develop facile method for quantifying the oligomerization kinetic of unstable protein solutions. One can also expect that our results enable to development of a procedure for preparing stable polymer particle conjugates with a controlled coverage of the hnRNPA LCD protein and the zeta potential.

## Author Contributions

PŻ: investigation (DLS and AFM), data curation, co-writing—original draft, PS, AK, ABS purified the protein, ABS calculated the theoretical charges of the proteins, PS the protein characterization in the bulk, stability of monolayers on latex particles performance of the experiments, data evaluation, method description, manuscript editing AM streaming potential performance and data curation investigation, cowriting—original draft, supervision, review and editing, formal analysis, ABS and ZA: conceptualization, writing – original draft, writing – review & editing, supervision, formal analysis funding acquisition, supervisions. All authors have read and agreed to the published version of the manuscript.

## Institutional Review Board Statement

Not applicable.

## Informed Consent Statement

Not applicable.

## Data Availability Statement

The data are available on request.

## Acknowledgments

hnRNPA2_LC_WT was a gift from Nicolas Fawzi (Addgene plasmid # 98657 ; http://n2t.net/addgene:98657; RRID:Addgene_98657). This work was financially supported by the National Science Center, Poland, Opus Project, 2021/43/B/ST8/01900.

