## Supplemental Figures for "An Effective Method for Determining the Degree of Oligomerization of hnRNPA2 Low Complexity Domain"

Supplementary Figures

**Figure 1S.** The amino acid sequences of human hnRNPA2 (UniProt: P22626). The studied sequence was from the amino acid 188 to 344, MW (LCD): 14607,91, theoretical pI (LCD): 9.45.

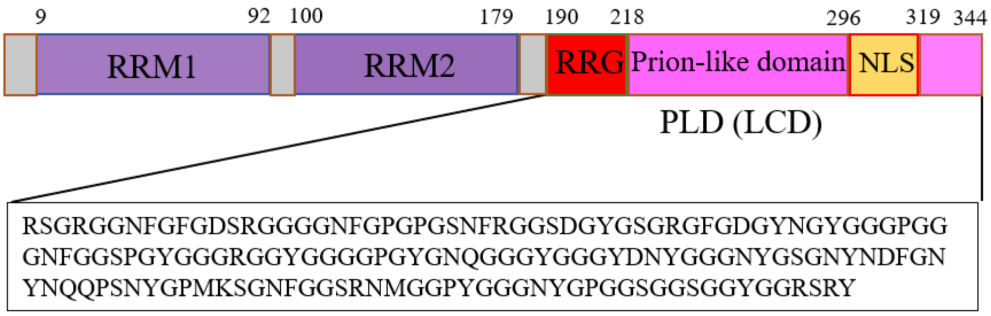

**Figure 2S.** SDS-PAGE representing purity of hnRNPA2 LCD after expression and purification from *E.coli* (band 14 represents the final protein solution).

1 2 3 4 5 6 7 8 9 10 11 12 13 14

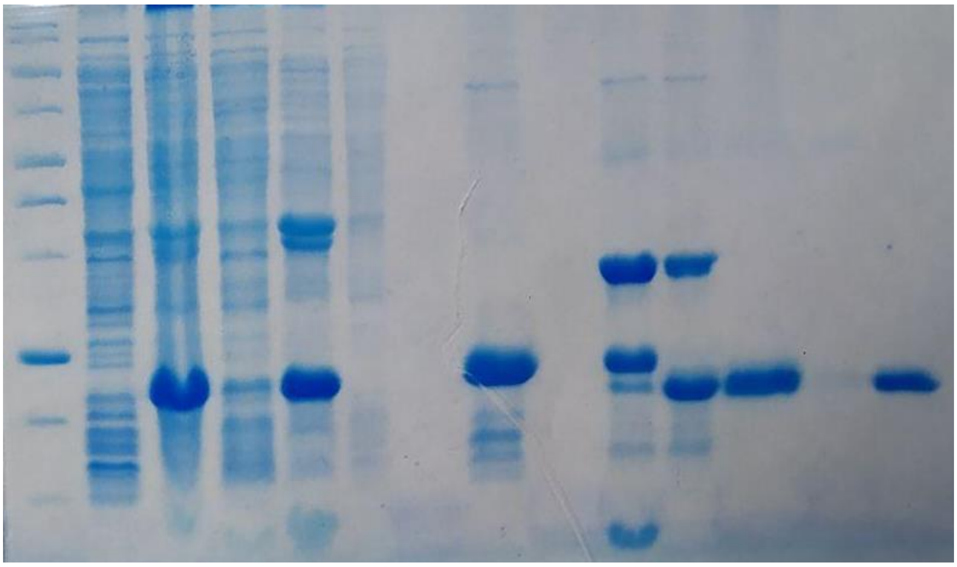

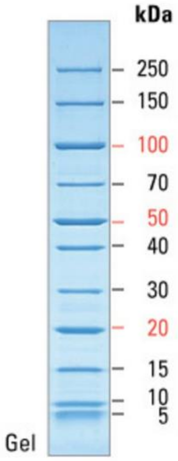


A description of the gel:

1 - standard mass PageRuler Unstained
2 - supernatant after first sonication
3 - pellet after first sonication
4 - supernatant after second sonication
5 - pellet after second sonication
6 - Flowthrough NiNTA
7 - Wash NiNTA (A6)
8 - Elution NiNTA
9 - A40 (desalting)
10 - A sample after desalting + TEV time "0"
11 - A sample after desalting + TEV overnight
12 - A sample after NiNTA 2
13 - A16 (NiNTA)
14 - SEC protein fraction A42

**Figure 3S.** hnRNPA2 Low Complexiy Domain with sodium ions structures determined by AlphaFold, [1-6], a) monomer (two different orientations), b) dimer (two different orientations), c) 13-mer (two different orientations).

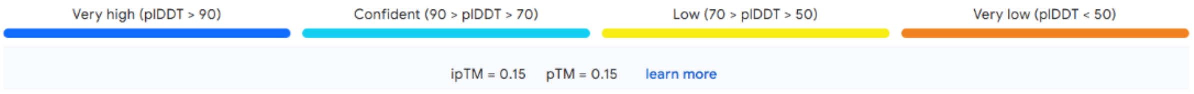


a) monomer b) dimer c) 13-mer

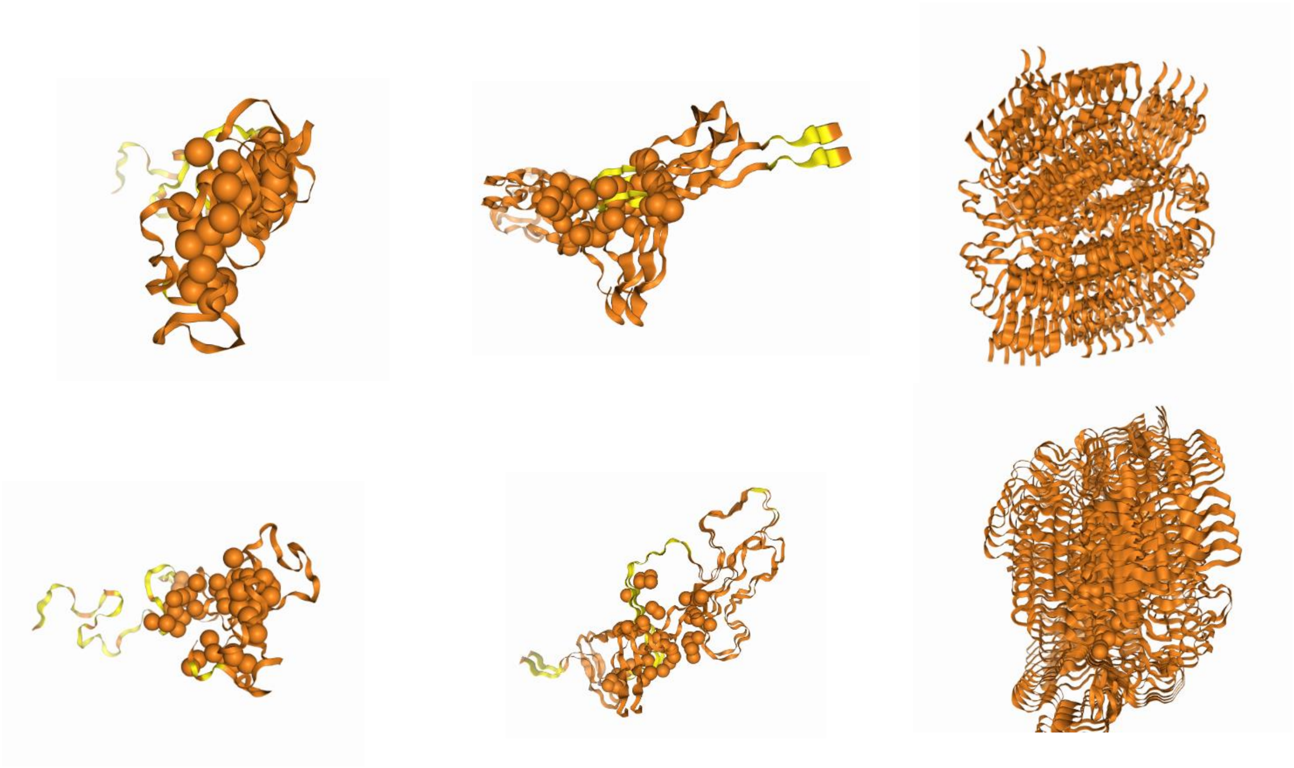

**Figure 4S.** AFM micrographs of hnRNPA2 LCD protein layers on mica Experimental conditions: bulk protein concentration $c_b$ = 0.5 mg $L^{-1}$, pH 4.0, 10 mM NaCl, adsorption time = 15 min, a) planar morphology of the mica covered by hnRNPA2 LCD, b) 3D AFM image of the mica covered by hnRNPA2 LCD.

a)

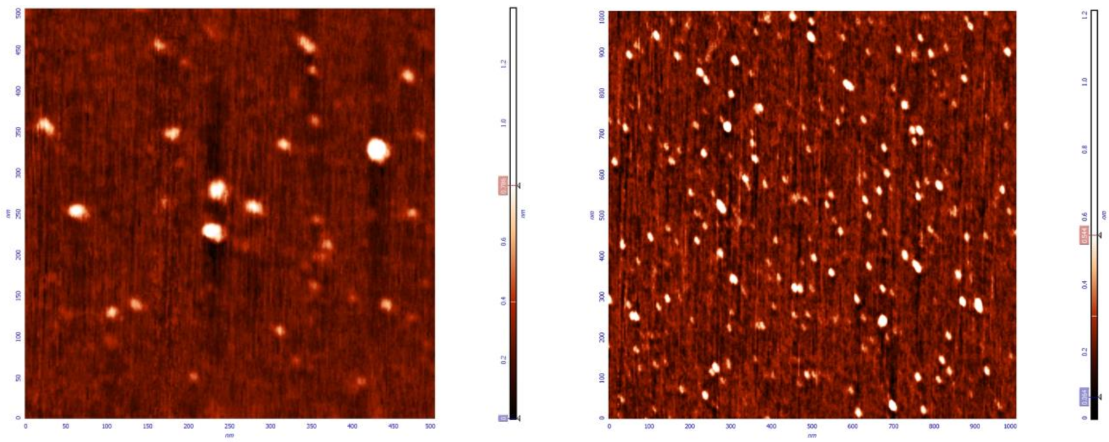

b)

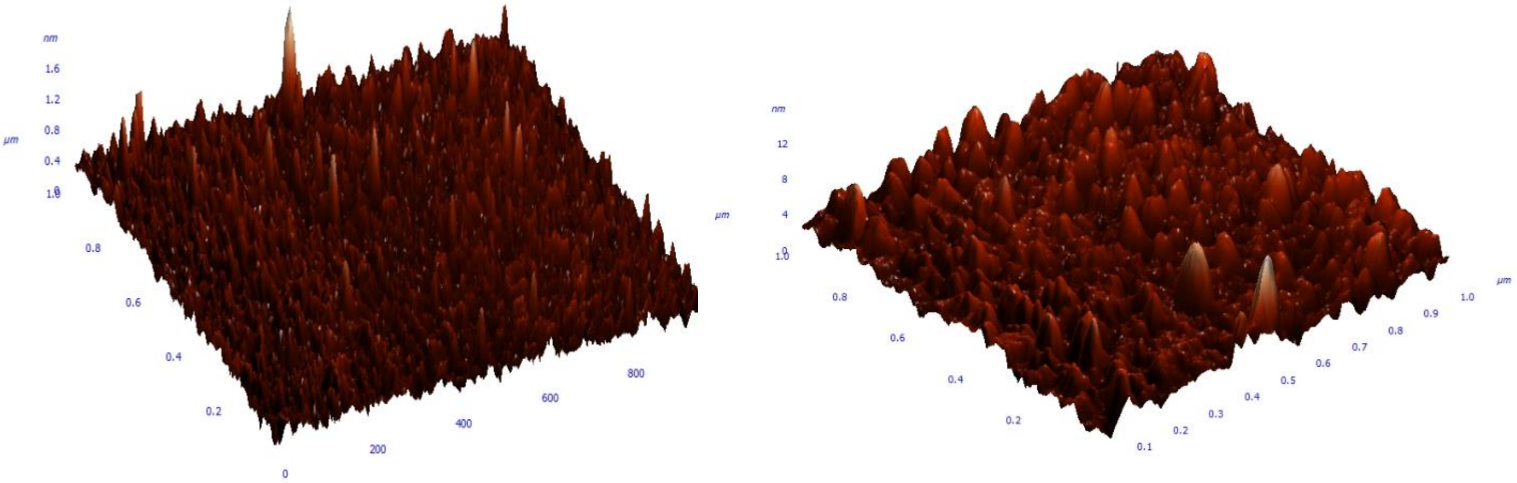

**Figure 5S.** Dependence of the zeta potential of the hnRNPA2 LCD layer on mica on the adsorption time acquired by SPM in the microfluidic cell (data points); volumetric flow rate of 0.02 mL $min^{-1}$, pH 4, 10 mM NaCl, protein bulk concentration 2.5 mg $L^{-1}$ (stock protein concentration 50 mg $L^{-1}$). The arrow shows the beginning of the desorption run, where pure electrolyte was flushed through the cell. The solid line shows theoretical results calculated using convective-diffusion and the electrokinetic models.

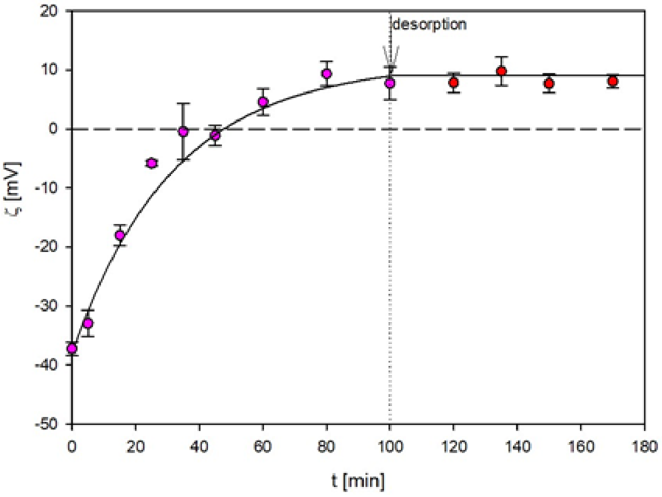


**Figure 6S.** The zeta potential of the hnRNPA2 LCD layer on mica vs pH acquired by the SPM in the microfluidic cell (10 mM NaCl). Filled points indicate the zeta potential for a discrete increase in pH from 4 to 9, whereas open points represent zeta potential for pH shift from 9 back to 4.

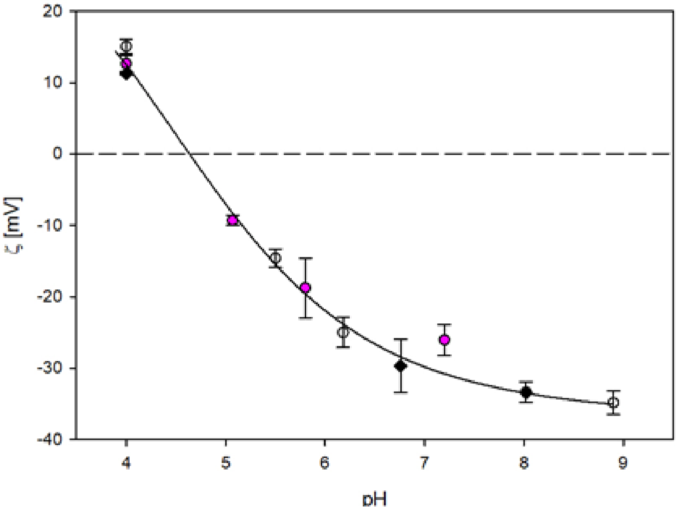
